# Chromosome-level Genome Assembly and Annotation of *Petunia hybrida*

**DOI:** 10.1101/2024.09.19.611905

**Authors:** Ali Saei, Donald Hunter, Elena Hilario, Charles David, Hilary Ireland, Azadeh Esfandiari, Ian King, Ella Grierson, Lei Wang, Murray Boase, Matthew Kramer, Shankar Shakya, Megan Bowman, Christopher R. Barbey, David Chagné

**Affiliations:** AgResearch Grasslands, Palmerston North, New Zealand; The New Zealand Institute for Plant and Food Research Limited (Plant & Food Research), Palmerston North, New Zealand; Plant & Food Research, Auckland, New Zealand; Plant & Food Research, Lincoln, New Zealand; Ball Horticultural Company, West Chicago, Illinois, USA

## Abstract

*Petunia hybrida* is the world’s most popular garden plant and is regarded as a supermodel for studying the biology associated with the Asterid clade, the largest of the two major groups of flowering plants. Unlike other Solanaceae, petunia has a base chromosome number of seven, not 12. This along with recombination suppression has previously hindered efforts to assemble its genome to chromosome level. Here we achieve a chromosome-level assembly for *P. hybrida* using a combination of short-read and long-read sequencing, optical mapping (Bionano) and Hi-C technologies. The resulting assembly spans 1253.6 Mb with a BUSCO score of 99.8%. A total of 35,089 genes were predicted and of those 29,655 were functionally annotated. Syntenic regions between petunia, tomato and pepper were identified, highlighting rearrangements that have occurred since their divergence indicating that the 12 chromosomes of Solanaceae did not originate from whole genome duplication of an ancestral species with seven chromosomes like petunia. This chromosome-level assembly will significantly enhance trait mapping efficiency in petunia and serve as a valuable resource for functional genomic studies in this key plant model.

## Introduction

Petunia (*Petunia* spp.) a genus first described by Jussieu in 1803, originates from South America and is globally recognized for its economic importance as an ornamental plant. Petunia is increasingly regarded as a model plant for the Asterid clade because of its numerous desirable traits ^[1]^. These traits include its short generation time, highly active endogenous transposon system, prolific seed production, and the ease with which it can be self-pollinated and crossed, propagated (sexually and asexually), transformed stably or transiently, and used with virus-induced-gene-silencing technology. The close evolutionary relationship of petunia with other members of the Solanaceae (e.g. potato, tomato, pepper, pepino and eggplant) makes it an excellent model for comparative genetics, particularly for studying traits related to evolutionary development biology, such as self-incompatibility, floral development, floral volatiles, petal senescence, inflorescence, and branching architecture ^[1]^. Commercial and standard laboratory lines of petunia are hybrid in origin, leading to their classification as *Petunia hybrida*. These hybrids have the same base chromosome number (2n = 2x = 14) as their origin species because of the species barriers being mainly pre-zygotic ^[1]^.

Whole-genome sequencing has been completed for several members of the Solanaceae family, including tomato ^[2]^, potato ^[3]^, *Physalis* ^[4]^, pepper ^[5]^, eggplant ^[6]^, tobacco ^[7]^ and pepino ^[8]^. Historically, the assembly of the petunia genome to chromosomal level has been hampered by both the lack of synteny with other Solanaceae (due to different chromosome number) and its low recombination frequency ^[9-11]^. Assembly of the two putative progenitors of *P. hybrida* was reported in 2016 by an international consortium ^[9]^, but the above-mentioned challenges and the limitations of sequencing technology at the time meant they were only able to resolve the *P. axillaris* and *P. inflata* genomes to 83,639 and 136,283 scaffolds, respectively. More recently the genome of *P. axillaris* has been resolved to seven chromosomes ^[12]^ and to 12 large scaffolds ^[13]^ by the University of Bern, and the genome of *P. hybrida* resolved to 1,226 scaffolds ^[14]^. Here we report on the *de novo* assembly of *P. hybrida* into its seven pseudochromosomes, followed by genome annotation. Comparative genome mapping was also performed against *P. axillaris* and other Solanaceae genomes, including tomato and pepper. We envisage that this high-quality genome will enable better utilisation of the functional analytical tools developed for petunia and contribute further to its status as a model for studying biology within the Asterid clade.

## Materials and Methods

### Plant material and growth conditions

Two inbred and closely related *Petunia hybrida* lines referred here as Line A and Line B were used for genome assembly. They were sourced from the breeding lines of Ball Horticultural Company, Chicago, USA. Vegetative clones of these inbred lines were used unless stated otherwise and were grown in individual pots in standard greenhouse facilities at Plant & Food Research, Palmerston North, New Zealand.

### Genomic DNA isolation and sequencing

We sequenced and assembled the genome of Line A using a combination of short-read sequencing, long-read sequencing, optical mapping and chromosome confirmation capture technology (Hi-C). High molecular weight genomic DNA (HMW gDNA) was isolated from ∼1 g of freshly harvested shoot tip tissue using an in-house CTAB/PVP/NaCl method that involved Proteinase K treatment of the isolated nuclei and a further chloroform: IAA extraction ^[15]^. Short read sequencing of the HMW gDNA was performed by the Australian Genome Research Facility (AGRF; Melbourne, Australia) with HiSeq HT chemistry and 125 bp paired-end (PE) sequencing (Illumina, San Diego, CA). Long-read sequencing of the HMW gDNA was carried out by the GrandOmics Biosciences Co., Ltd, (Beijing, China) using single-molecule real-time (SMRT) sequencing on the PacBio Sequel platform (Pacific Biosciences, Menlo Park, CA). Single-molecule optical maps were generated and analysed at Kansas State University with the Irys® genome mapping system (Bionano Genomics, CA) from the HMW gDNA of >100 Mb in size that had been isolated in agarose plugs by the method described in ^[16]^. Hi-C libraries were constructed from plant tissue using the Phase Genomics Proximo™ Hi-C kit Plant Protocol Kit Version 1 (Phase Genomics, Seattle, Washington, USA) and the gDNA sample was double size selected (0.6/0.2X) to obtain an amplicon average fragment size of 480 bp. The sample was sequenced at AGRF to obtain 150 bp PE reads with the Illumina MiSeq platform.

### Genome assembly and scaffolding

The reads were first error corrected using Canu v1.7 software ^[17]^. The corrected reads were then used for genome size estimation by generating a 33-mer frequency distribution with Jellyfish v2.3.0 and the results analysed with GenomeScope v1 with maximum k-mer coverage set to –1 ^[18]^. Finally, the corrected reads were trimmed and assembled using Canu with correctedErrorRate = 0.035 and corMhapSensitivity = normal. The assembly was polished with high quality Illumina PE reads using Pilon ^[19]^. Completeness of the genome was assessed using the Benchmarking Universal Single Copy Ortholog (BUSCO) tool version 5.6.1 using the viridiplantae_odb10 and solanales_odb10 databases ^[20]^. The Irys® generated single molecule maps were assembled de novo using the Bionano Solve v3.1.0 assembly pipeline. The assembled optical maps were used to order, orientate, validate, and scaffold the Canu assembly. This was achieved by digesting the PacBio assembly in silico with the same restriction enzymes used for generating the single molecule optical maps and aligning the PacBio and optical maps using the Bionano Solve v3.1.0 Hybrid Scaffolding pipeline. A Hi-C ^[21]^ scaffolding pipeline developed in-house at Plant & Food Research was further used for validating, ordering and orientating the hybrid PacBio-Bionano scaffolds. The utility of this in-house Hi-C method was demonstrated by it having successfully scaffolded two genomes for which confirmation by synteny was available ^[22,23]^. We used bwa mem ^[24]^ to align Hi-C reads to the PacBio-Bionano scaffolds, using specific options (−5, -S, and - P) for optimization as detailed in bwa mem usage. The -5 option reduces the amount of secondary and alternate mappings, addressing Hi-C data ambiguity. Simultaneously, the -S and -P options enhance alignment efficiency by avoiding assumptions about reads typical in other library types. Low-quality mapped reads were filtered out and remaining reads binned with 20 kb bin length to generate Hi-C contact data. The contact data were normalized using ICE ^[25]^. The *P. axillaris* datasets used for genomic structural comparison were obtained from NCBI PRJNA689605 ^[12]^, PRJNA858035 ^[13]^.

### RNA isolation, library construction, and transcriptome sequencing

Line B plants were grown from seed in individual pots in the Palmerston North greenhouse facility for transcriptome sequencing of their floral limb tissue at three developmental stages (young bud, half open and full open). Limb tissue was collected from these mixed stage flowers for sequencing. Total RNA was isolated from the limb samples using the Qiagen RNAeasy Plant Minikit according to manufacturer’s instructions. RNA concentration was determined using a Nanodrop1000 spectrophotometer (ThermoFisher Scientific). RNA quality was determined with a Fragment Analyser (Advanced Analytical Technologies, Inc.) using RNA (Eukaryotic) Analysis mode of the PROSize 2.0 software. Libraries were constructed using a modification of the method of ^[26]^. Indexed 25 μl samples of concentrations > 17 nM and compatible with single pooling were sent to the Australian Genome Research Ltd. (AGRF) for 100 bp PE sequencing.

### Identifying repeat sequences and genome annotation

RepeatModeler v2.0.4 ^[27]^ was used to perform *de novo* identification of repetitive elements and generate repeat libraries for subsequent repeat masking within the assembled genome. Following this, RepeatMasker v4.1.5 (www.repeatmasker.org) was applied for the annotation and masking of identified repeats, using the repeat library generated by RepeatModeler. The Extensive de-novo TE Annotator (EDTA v2.1.0) ^[28]^ was used for the identification and classification of transposable elements.

The genome annotation was performed using the Braker3 pipeline v3.0.6, a comprehensive tool for structural and functional genome annotation ^[29]^. The input for the pipeline included the masked genome and a diverse set of protein sequence references and transcriptome data. These protein references included the SwissProt (release 2024_03), the Kyoto Encyclopedia of Genes and Genomes (KEGG; release 110), The Arabidopsis Information Resource 10 (TAIR10; release 20230630), and the hierarchical catalogue of orthologs (OrthoDB v11) Viridiplantae databases. Protein sequences from a previous petunia genome (*P. axillaris* v1.6.2) ^[9]^ were used to capture lineage-specific information and improve annotation precision.

Gene prediction was performed using Augustus v3.4.0, a highly accurate *ab initio* gene prediction tool that leverages hidden Markov models ^[30]^. Functional annotation of predicted genes was conducted using EggnogMapper v2.1.12 ^[31]^, which assigns orthology and functional annotations based on the EggNOG database (v5.0). To process and manipulate the annotation files, gffread v0.12.7 and Another Gtf/Gff Analysis Toolkit (AGAT v1.4.0) were utilized. These tools facilitated the conversion, filtering, and validation of GFF files, ensuring the integrity and consistency of the annotation data. The output files generated from the annotation process were a GFF3 file, a protein FASTA file, and a set of EggnogMapper annotations.

### Synteny analysis

Syntenic blocks between petunia and other Solanaceae were produced using “nucmer –mum”, then “delta-filter” with 70% nucleotide identity (-i) and syntenic block length of 1 kb (-l) settings for every comparison ^[32]^. Syntenic blocks were defined as regions containing at least 20 orthologous loci between both genomes. Syntenic blocks between *P. hybrida* and *P. axillaris* were produced using “nucmer –mum”, then “delta-filter” with 90% nucleotide identity (-i) and syntenic block length of 2 kb (-l) settings for every comparison. Circos plots for genomic synteny between petunia and other Solanaceae were generated using the Circos visualisation tool (v0.69-6). Linear synteny plots were produced using pyGenomeViz[33].

## Results

### PacBio long-read sequencing and assembly

Approximately 16.5 million raw PacBio long-reads were generated with an average length of 8.9 kb and an N50 of 12.8 kb. The genome size of *P. hybrida* was estimated to be 1.15 Gb based on 33-mer analysis (Supplementary Figure S1). The PacBio data provided approximately 127-fold coverage of the estimated genome size. Canu assembled the PacBio data into 781 contigs for a total assembly size of 1,267,612,960 bp with a contig length N50 of 7.1 Mb (Table 1, Supplementary Figure S2). The contigs were then polished with 82 Gb of short-read sequences (∼80-fold coverage).

**Table 1.** Summary statistics of the *Petunia hybrida* genome assembly.

| Metric | Pacbio scaffolds | Hybrid Pacbio-Bionano scaffolds | Hi-C scaffolds |
| --- | --- | --- | --- |
| Assembly Size (bp) | 1,267,612,960 | 1,250,888,111 | 1,267,083,507 |
| N (%) | 0 | 0.16 | 0.16 |
| GC (%) | 38.3 | 38.29 | 37.81 |
| AT (%) | 61.7 | 61.71 | 62.19 |
| scaffold count | 781 | 184 | 7 |
| longest scaffold (bp) | 27,118,569 | 39,350,372 | 227,137,022 |
| scaffold N50 length (bp) | 7,069,514 | 10,897,467 | 174,246,978 |
| scaffold N50 count | 54 | 36 | 4 |
| scaffold N90 length (bp) | 2,273,843 | 3,677,139 | 162,799,232 |
| scaffold N90 count | 169 | 117 | 7 |
| contig count | 781 | 548 | 26,268 |
| contig N50 length (bp) | 7,069,514 | 6,993,002 | 7,084,997 |
| contig N50 count | 54 | 55 | 54 |
| contig N90 length (bp) | 2,273,843 | 2,389,267 | 2,420,040 |
| contig N90 count | 169 | 169 | 1 |

### Genome scaffolding using optical mapping

Optical mapping generated 176.9 Gb of single molecule fragments each greater than 150 kb in length and containing a minimum of nine stainable sites. The N50 molecule length (≥150 kb) was 225 kb, meaning that at least half of the genome was covered by molecules larger than 225 kb on average after discarding molecules smaller than 150 kb. This provided ∼153-fold coverage of the estimated genome size. The single molecule fragments were assembled *de novo* into 2701 consensus genome maps with N50 of 520 kb that covered 1.27 Gb of the total genome map length. The consensus maps were then used to order, orientate, validate, and scaffold the PacBio contigs digested similarly, but *in silico*. A total of 329 out of the 781 PacBio contigs were ordered and oriented into 184 hybrid scaffolds with N50 length of 10.9 Mb, which covered ∼99% of the assembled genome (Table 1). An example of how the optical mapping data ordered, oriented, scaffolded and independently validated the PacBio data is illustrated in Supplementary Figure S3.

### Anchoring of hybrid scaffolds with Hi-C

Approximately 1.94 billion 150 bp Hi-C reads were used to anchor and orientate the hybrid PacBio-Bionano scaffolds into a chromosomal level assembly. In total, 184 hybrid PacBio-Bionano scaffolds and a further 24 PacBio contigs were anchored by Hi-C into seven pseudochromosomes with total genome size of 1,267,083,507 bp (Figure 1; Table 1). The lengths of each pseudochromosome ranged from 160.7 to 224.3 Mb (Table 2). Their numbers were assigned based on alignment with published linkage maps of petunia species ^[9]^ and a genome of *P. axillaris* genome resolved to its seven chromosomes ^[12]^. The centromeric region of each chromosome was clearly identified in the Hi-C data. These regions showed strong interactions both within themselves and with other centromeric regions, while displaying weak interactions with non-centromeric regions (Figure 1). There was considerable structural similarity between the *P. hybrida* genome and the *P. axillaris* genome (Figure 2) validating the overall assembly of both genomes. Unexpectedly, synteny analysis between the *P. hybrida* and *P. axillaris* genomes identified a notable difference between both assemblies, with both end regions of chromosome 1 (∼ 28 Mb at beginning and ∼ 22 Mb at end) being present at the opposite ends in the *P. axillaris* assembly. However, the *P. hybrida* chromosome 1 assembly was validated as correct by a perfect collinearity of that chromosome with a separate *P. axillaris* genome assembly ^[13]^ (Figure 3A). Moreover, the Hi-C contact map of *P. hybrida* chromosome 1 did not support any miss-assemblies in the two putative regions of *P. hybrida* chromosome 1 where the breakpoints would have been located (Figure 3B and C).

**Table 2.**
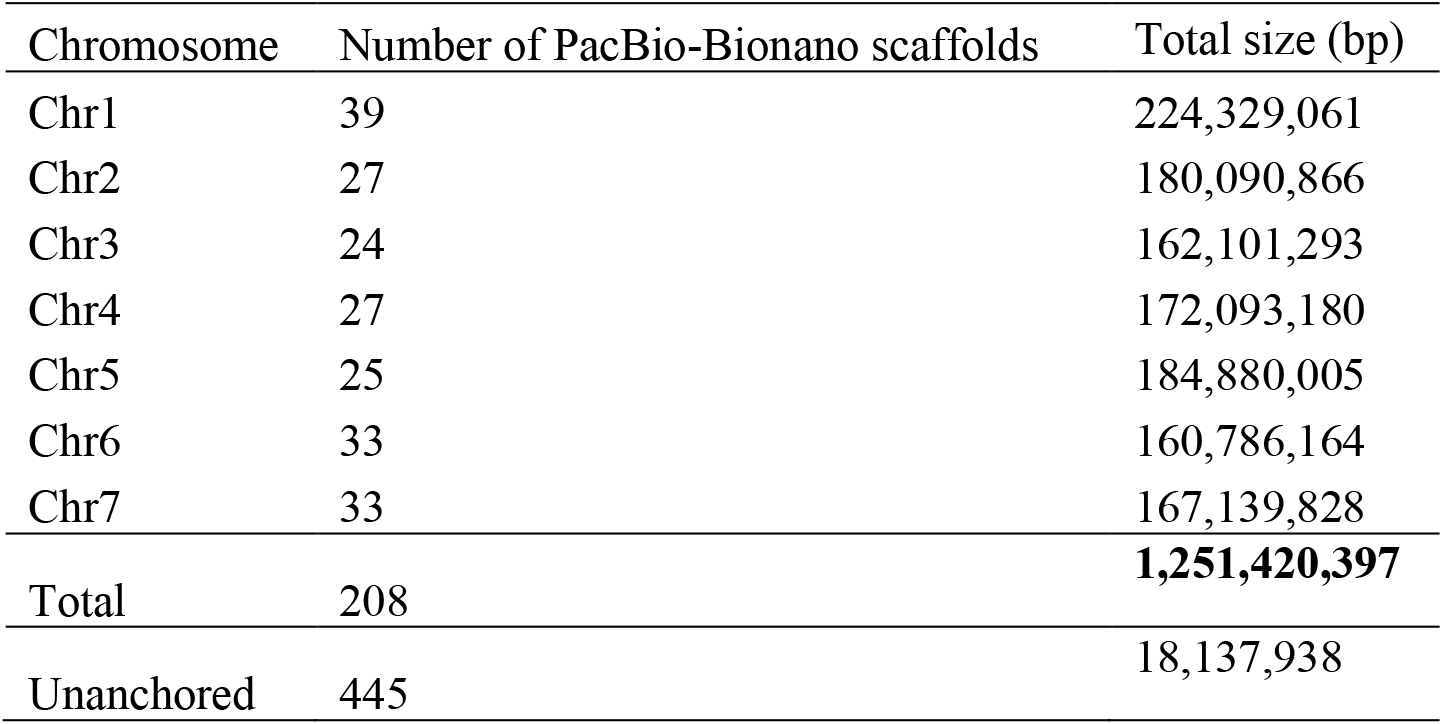
Summary statistics of the seven pseudochromosomes of the *Petunia hybrida* genome assembly.

**Figure 1.**
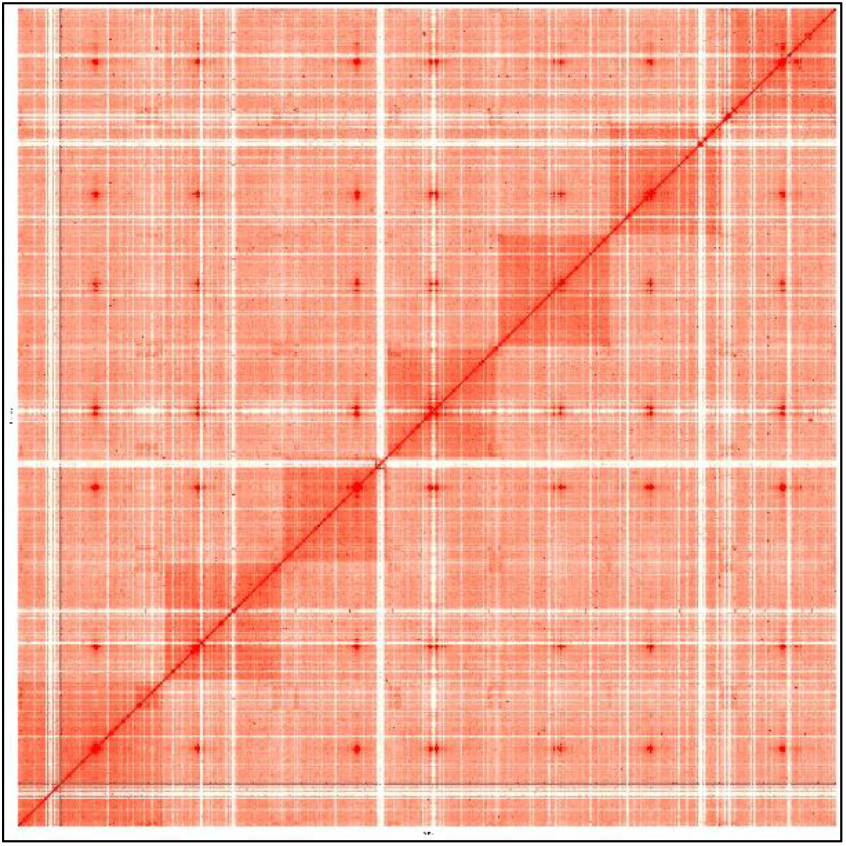
Hi-C contact map of the *Petunia hybrida* genome. The plot was generated with a 300 kb bin size. The map shows the calculated interaction frequency distribution of Hi-C links both between and within chromosomes. The heatmap colors (light red to dark red) indicate the frequency of Hi-C interaction links, ranging from low to high. The single dark red region within each chromosome is the centromere.

**Figure 2.**
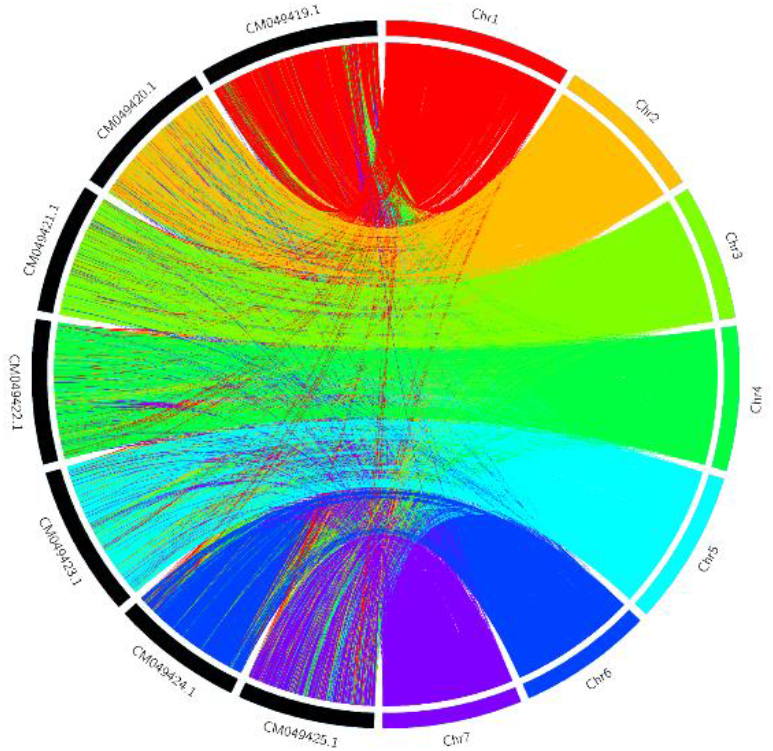
Circos plot illustrating the structural similarities between *Petunia hybrida* and *Petunia axillaris* genomes.

**Figure 3.**
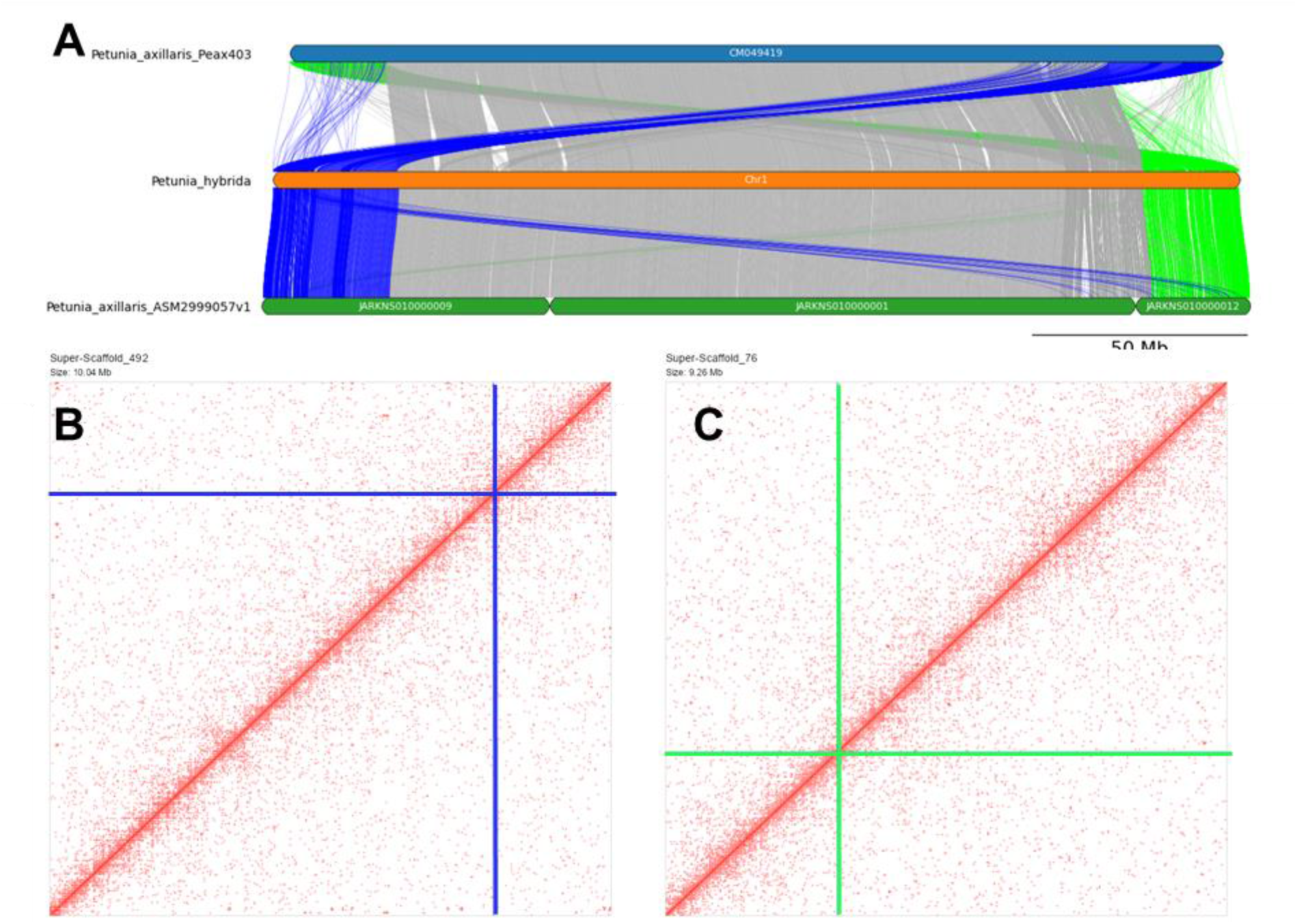
Confirmation of *Petunia hybrida* chromosome 1 assembly. A. Synteny plot showing discrepancy between the assembly of *P. hybrida* (middle track) and *P. axillaris* (top track, PRJNA689605 ^[12]^) and concordance with a separate independently scaffolded *P. axillaris* assembly (bottom track, PRJNA858035 ^[13]^). B, and C, Hi-C plots, marked by the blue and green crosshairs, highlight the junctions in the *P. hybrida* scaffolds where the corresponding breaks were in the *P. axillaris* assembly.

BUSCO analysis of the final assembly using the viridiplantae_odb10 dataset revealed that 99.8% BUSCOs (single: 98.4%, duplicated: 1.2%, fragmented: 0.2%, missing: 0.0%, n: 425) were present.

This was 97.6% when the solanales_odb10 dataset comprising 5950 genes was used in the BUSCO analysis (Supplementary Figure S4).

### Genome annotation

In total, approximately 894 Mb (70.46%) of the assembled petunia genome comprised interspersed repeats (Table 3). Of these sequences, 347,312 repeats were retroelements of which 310,112 were long-terminal repeats (LTR) spanning 482 Mb (38% of the genome). These included 73,447 Ty1/Copia and 221,564 Gypsy/DIRS1 retroelements spanning 96 Mb (7.5%) and 369 Mb (29%) respectively of the petunia genome. A total of 35,089 genes were predicted and of those 29,655 were functionally annotated.

**Table 3.** Repeated elements analysis in the *Petunia hybrida* genome assembly.

| Repeat type | Number of elements | Length (bp) | Percentage of sequence (%) |
| --- | --- | --- | --- |
| Retroelements | 347,312 | 508,500,044 | 40.05 |
| **SINEs:** | **3,457** | **412,955** | **0.03** |
| **Penelope:** | **184** | **39,515** | **0.00** |
| **LINEs:** | **33,743** | **26,224,252** | **2.07** |
| CRE/SLACS | 0 | 0 | 0.00 |
| L2/CR1/Rex | 0 | 0 | 0.00 |
| R1/LOA/Jockey | 0 | 0 | 0.00 |
| R2/R4/NeSL | 0 | 0 | 0.00 |
| RTE/Bov-B | 9,817 | 5,263,192 | 0.41 |
| L1/CIN4 | 22,334 | 20,411,414 | 1.61 |
| **LTR elements:** | **310,112** | **481,862,837** | **37.95** |
| BEL/Pao | 0 | 0 | 0.00 |
| Ty1/Copia | 73,447 | 96,100,243 | 7.57 |
| Gypsy/DIRS1 | 221,564 | 369,159,465 | 29.08 |
| Retroviral | 1,295 | 660,318 | 0.05 |
| **DNA transposons** | **49,309** | **37,375,725** | **2.94** |
| hobo-Activator | 5,808 | 5,871,997 | 0.46 |
| Tc1-IS630-Pogo | 0 | 0 | 0.00 |
| En-Spm | 0 | 0 | 0.00 |
| MULE-MuDR | 34,241 | 22,413,695 | 1.77 |
| PiggyBac | 0 | 0 | 0.00 |
| Tourist/Harbinger | 668 | 259,146 | 0.02 |
| **Rolling-circles** | **5,258** | **3,006,527** | **0.24** |
| **Unclassified:** | **1,062,399** | **348,633,296** | **27.46** |
| **Total interspersed repeats:** |  | **894,548,580** | **70.46** |

### Synteny between petunia and other Solanaceae

Comparative genome mapping was performed between petunia and the published Solanaceae genomes, tomato, potato, *Physalis*, pepper and tobacco (Figure 4; Supplementary Figure S5). Synteny between petunia and the other Solanaceae is fragmented into small syntenic blocks in multiple chromosomes rather than broad chromosome-scale fusions (Figure 4; Supplementary Figure S5). Pairwise comparison between petunia and tomato, and petunia and pepper (Figure 4A), identified 6,145 and 6,691 orthologous loci of sequence similarity greater than 70% and longer than 1,000 bp, respectively (Supplementary Figure S5). Of these, 5,768 and 5,724 loci defined 37 and 43 syntenic blocks made of more than 20 colinear orthologous loci between petunia and tomato, and between petunia and pepper, respectively. Visualisation of alignments using linear syntenic plots indicated that orthologous loci and syntenic blocks hit a single region in tomato and pepper (Figure 4B), indicating that the 12 chromosomes of Solanaceae did not come from whole genome duplication of an ancestral species with seven chromosomes like petunia. Syntenic blocks tended to be in pericentric regions.

**Figure 4.**
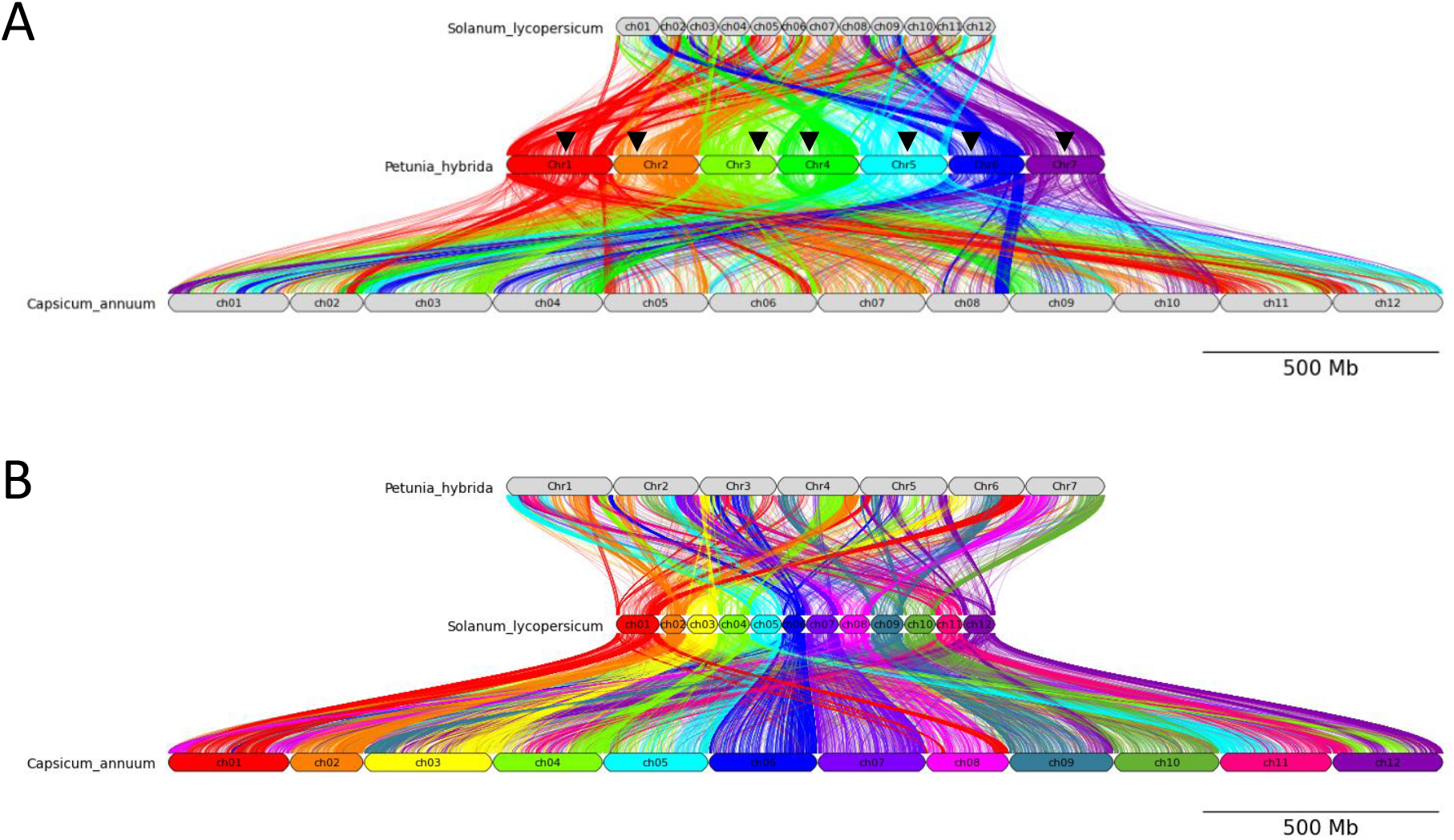
Conserved synteny across petunia, tomato and pepper. Each line represents a region longer than 1 kb with a sequence similarity of 70% or higher. A) Colours correspond to chromosomes of petunia (*Petunia hybrida)*. B) Colours correspond to chromosomes of tomato (*Solanum lycopersicum)* and pepper *(Capsicum annum)*. Centromere position in each *P. hybrida* chromosome is indicated by black arrowhead.

## Discussion

### A near-complete assembly of petunia

We have constructed the first near-complete genome assembly for *P. hybrida*. We achieved chromosome-scale scaffolding that has synteny with the genomes of other Solanaceae species. This assembly incudes seven pseudochromosomes covering a total of 1,253.6 Mb and a BUSCO completeness of 99.6%, which is a vast improvement when compared with the highly fragmented petunia genomes published previously ^[9]^. The high contiguity and completeness were made possible through combination of long-read PacBio sequencing, optical mapping (Bionano) and Hi-C technologies. These complementary techniques, focusing on different but overlapping scales allowed us to produce a high-quality genome. In addition to ordering, orienting, and scaffolding contigs, the Bionano maps provided an extra layer of validation to ensure the accuracy of the PacBio contig assembly. Using such a hybrid assembly approach was crucial for identifying and resolving mis-assemblies, such as chimeric joins, inversions, and translocations present in the initial PacBio contigs. The Hi-C method was able to anchor the hybrid PacBio-Bionano scaffolds to pseudochromosomes.

Hi-C was advantageous for scaffolding petunia genome as high-density linkage maps can be challenging to obtain due to hybrid species such as *P. hybrida* having extensive differences in recombination rate. This recombination suppression in petunia makes building linkage maps a very challenging task, which prevents the use of linkage maps for the purpose of anchoring genomes. Hi-C provided an ideal alternative for that purpose.

The combination of sub-tools within the Braker3 pipeline facilitated a comprehensive and accurate annotation of the genome, yielding high-confidence gene predictions and functional annotations that are crucial for downstream analyses and biological interpretation. Repeat annotation of the new petunia genome identified a total of 894 Mb repeated elements (70.5% of the genome). The number of repeats and genome size covered by them is larger than the previous petunia genome assemblies ^[9]^, indicating that our long reads-based assembly is more accurate at resolving these genome features.

Among the repeated sequences, 481 Mb were LTRs. The tomato genome has 401 Mb covered by LTRs ^[2]^, which means that such classes of repeated elements cannot alone explain the size difference between tomato and petunia genomes. It is possible that the unclassified portion of the repeated elements spanning 348 Mb may explain the larger size of the petunia genome.

### Synteny between petunia and Solanaceae

Our chromosome-level assembly of *P. hybrida* enabled whole-genome comparative mapping between petunia and other Solanaceae. Although conserved synteny was observed across genomic regions, our results suggest multiple chromosomal rearrangements since the divergence of these species.

Chromosomal rearrangements have been reported in the Solanaceae previously ^[34-37]^, though most comparisons involved genomes with 12 chromosomes. The new *P. hybrida* genome allowed for synteny analysis between species with seven and 12 chromosomes. Our comparative genome mapping and evidence for macro-syntenic regions and rearrangements between petunia and tomato suggests that some putative chromosomal fusions occurred during species radiation. More research, including genome sequencing of Solanaceae species with varying numbers of chromosomes (e.g. *Calibrachoa* n = 9), combined with cytological studies will be required to fully understand how these rearrangements contributed to speciation within the Solanaceae family.

## Conclusion

We have constructed a near-complete genome assembly of *Petunia hybrida*, which serves as a valuable resource for future functional genomic research in this important model plant. The pseudo-chromosome assembly will facilitate more efficient trait mapping in petunia and provide a reference for studying recombination rate variability within the genus. While macro-synteny exists between petunia and tomato, several chromosomal rearrangements, including fusions, have occurred. Further synteny analysis and cytological studies are needed to unravel genome evolution within the Solanaceae family

## Authors contributions

AS, DH and DC designed the study and wrote the manuscript. AS assembled the genome. EH isolated high molecular weight DNA. EG and LW isolated RNA. HI produced the RNA-seq libraries. IK and AE managed clonal plant maintenance. AS, CD, SS, MB and CRB did repeat and gene annotations. DH and MK initiated the project. MK supplied the study materials.

## Acknowledgements

Funding for this project was provided by Ball Horticulture Company, Chicago, USA. DC and EH received funding from the New Zealand MBIE SSIF platform Genomics Aotearoa. We thank Sarah Bailey, Rebecca Bloomer and Ed Morgan (Plant & Food Research) for their comments on the manuscript.

## Data availability

The genome assembly of *Petunia hybrida* reported here has been submitted to NCBI under BioProject PRJNA1156804 and BioSample accession SAMN43501245. The assembly and annotation files have also been deposited on FigShare ^[38]^ and SolGenomics.

## Conflict of interest

The authors declare no conflict of interest.

## Notes

https://doi.org/10.6084/m9.figshare.26940946

### Competing Interest Statement

The authors have declared no competing interest.

## References

1. Vandenbussche M, Chambrier P, Bento SR, Morel P. 2016. Petunia, your next supermodel? Frontiers in Plant Science, 7.

2. Sato S, Tabata S, Hirakawa H, Asamizu E, Shirasawa K, Isobe S, Kaneko T, Nakamura Y, Shibata D, Aoki K et al. 2012. The tomato genome sequence provides insights into fleshy fruit evolution. Nature, 485(7400):635–641.

3. Xu X, Pan SK, Cheng SF, Zhang B, Mu DS, Ni PX, Zhang GY, Yang S, Li RQ, Wang J et al. 2011. Genome sequence and analysis of the tuber crop potato. Nature, 475(7355):189–U194.

4. Lu, J, Luo, M, Wang, L et al. The Physalis floridana genome provides insights into the biochemical and morphological evolution of Physalis fruits. Horticulture Research 8, 244 (2021). 10.1038/s41438-021-00705-w

5. Qin C, Yu CS, Shen YO, Fang XD, Chen L, Min JM, Cheng JW, Zhao SC, Xu M, Luo Y et al. 2014. Whole-genome sequencing of cultivated and wild peppers provides insights into Capsicum domestication and specialization. Proceedings of the National Academy of Sciences of the United States of America, 111(14):5135–5140.

6. Hirakawa H, Shirasawa K, Miyatake K, Nunome T, Negoro S, Ohyama A, Yamaguchi H, Sato S, Isobe S, Tabata S et al. 2014. Draft genome sequence of eggplant (Solanum melongena L.): the representative Solanum species indigenous to the old world. DNA Research, 21(6):649–660.

7. Ko S-R, Lee S, Koo H, Seo H, Yu J, Kim Y-M, Kwon S-Y, Shin A-Y. 2024. High-quality chromosome-level genome assembly of Nicotiana benthamiana. Scientific Data, 11(1):386.

8. Song X, Liu H, Shen S, Huang Z, Yu T, Liu Z, Yang Q, Wu T, Feng S, Zhang Y et al. 2022. Chromosome-level pepino genome provides insights into genome evolution and anthocyanin biosynthesis in Solanaceae. The Plant Journal, 110(4):1128–1143.

9. Bombarely A, Moser M, Amrad A, Bao M, Bapaume L, Barry CS, Bliek M, Boersma MR, Borghi L, Bruggmann R et al. 2016. Insight into the evolution of the Solanaceae from the parental genomes of Petunia hybrida. Nature Plants, 2(6).

10. Bossolini E, Klahre U, Brandenburg A, Reinhardt D, Kuhlemeier C. 2011. High resolution linkage maps of the model organism Petunia reveal substantial synteny decay with the related, genome of tomato. Genome, 54(4):327–340.

11. Guo YF, Lin WK, Chen QX, Vallejo VA, Warner RM. 2017. Genetic determinants of crop timing and quality traits in two interspecific Petunia recombinant inbred line populations. Scientific Reports, 7.

12. National Center for Biotechnology Information (NCBI). PRJNA689605 Petunia axillaris subsp. axillaris cultivar:N Genome sequencing and assembly. University of Bern, 2022-12-16, https://ftp.ncbi.nlm.nih.gov/genomes/genbank/plant/Petunia_x_axillaris/

13. National Center for Biotechnology Information (NCBI). PRJNA858035 Petunia axillaris subsp. axillaris cultivar:P University of Bern, 2023-05-12 https://ftp.ncbi.nlm.nih.gov/genomes/genbank/plant/Petunia_axillaris/

14. National Center for Biotechnology Information (NCBI). PRJNA1060430 Petunia x hybrida Genome sequencing and assembly . 2024-01-02 https://ftp.ncbi.nlm.nih.gov/genomes/genbank/plant/Petunia_x_hybrida/

15. Hilario E: Plant nuclear genomic DNA preps. In. protocols.io; 2018.

16. Hilario E: Optical mapping preps for Petunia spp. In. protocols.io; 2019.

17. Koren S, Walenz BP, Berlin K, Miller JR, Bergman NH, Phillippy AM. 2017. Canu: scalable and accurate long-read assembly via adaptive k-mer weighting and repeat separation. Genome Research, 27(5):722–736.

18. Vurture GW, Sedlazeck FJ, Nattestad M, Underwood CJ, Fang H, Gurtowski J, Schatz MC. 2017. GenomeScope: fast reference-free genome profiling from short reads. Bioinformatics, 33(14):2202–2204.

19. Walker BJ, Abeel T, Shea T, Priest M, Abouelliel A, Sakthikumar S, Cuomo CA, Zeng QD, Wortman J, Young SK et al. 2014. Pilon: An integrated tool for comprehensive microbial variant detection and genome assembly improvement. Plos One, 9(11).

20. Waterhouse RM, Seppey M, Simao FA, Manni M, Ioannidis P, Klioutchnikov G, Kriventseva EV, Zdobnov EM. 2018. BUSCO applications from quality assessments to gene prediction and phylogenomics. Molecular Biology and Evolution, 35(3):543–548.

21. Burton JN, Adey A, Patwardhan RP, Qiu RL, Kitzman JO, Shendure J. 2013. Chromosome-scale scaffolding of de novo genome assemblies based on chromatin interactions. Nature Biotechnology, 31(12):1119-+.

22. Ireland HS, Wu C, Deng CH, Hilario E, Saei A, Erasmuson S, Crowhurst RN, David KM, Schaffer RJ, Chagné D. 2021. The Gillenia trifoliata genome reveals dynamics correlated with growth and reproduction in Rosaceae. Horticulture Research, 8(1).

23. Wu C, Deng C, Hilario E, Albert NW, Lafferty D, Grierson ERP, Plunkett BJ, Elborough C, Saei A, Günther CS et al. 2022. A chromosome-scale assembly of the bilberry genome identifies a complex locus controlling berry anthocyanin composition. Molecular Ecology Resources, 22(1):345–360.

24. Li H, Durbin R. 2009. Fast and accurate short read alignment with Burrows-Wheeler transform. Bioinformatics, 25(14):1754–1760.

25. Imakaev M, Fudenberg G, McCord RP, Naumova N, Goloborodko A, Lajoie BR, Dekker J, Mirny LA. 2012. Iterative correction of Hi-C data reveals hallmarks of chromosome organization. Nature Methods, 9(10):999-+.

26. Zheng H, Yin J, Gao Z, Huang H, Ji X, Dou C. 2011. Disruption of Chlorella vulgaris cells for the release of biodiesel-producing lipids: a comparison of grinding, ultrasonication, bead milling, enzymatic lysis, and microwaves. Appl Biochem Biotechnol, 164(7):1215–1224.

27. Flynn JM, Hubley R, Goubert C, Rosen J, Clark AG, Feschotte C, Smit AF. 2020. RepeatModeler2 for automated genomic discovery of transposable element families. Proceedings of the National Academy of Sciences of the United States of America, 117(17):9451–9457.

28. Su W, Ou S, Hufford MB, Peterson T: A tutorial of EDTA: extensive de novo TE annotator 2021. In: Plant Transposable Elements: Methods and Protocols. Edited by Cho J. New York, NY: Springer US: 55–67.

29. Hoff KJ, Lange S, Lomsadze A, Borodovsky M, Stanke M. 2016. BRAKER1: unsupervised RNA-Seq-based genome annotation with GeneMark-ET and AUGUSTUS. Bioinformatics, 32(5):767–769.

30. Stanke M, Schöffmann O, Morgenstern B, Waack S. 2006. Gene prediction in eukaryotes with a generalized hidden Markov model that uses hints from external sources. Bmc Bioinformatics, 7.

31. Huerta-Cepas J, Forslund K, Coelho LP, Szklarczyk D, Jensen LJ, von Mering C, Bork P. 2017. Fast genome-wide functional annotation through orthology assignment by eggNOG-Mapper. Molecular Biology and Evolution, 34(8):2115–2122.

32. Marçais G, Delcher AL, Phillippy AM, Coston R, Salzberg SL, Zimin A. 2018. MUMmer4: A fast and versatile genome alignment system. PLOS Computational Biology, 14(1):e1005944.

33. Shimoyama, Y. 2022. pyGenomeViz: A genome visualization python package for comparative genomics. https://github.com/moshi4/pyGenomeViz

34. Wu F, Tanksley SD. 2010. Chromosomal evolution in the plant family Solanaceae. BMC Genomics, 11(1):182.

35. Tanksley SD, Bernatzky R, Lapitan NL, Prince JP. 1988. Conservation of gene repertoire but not gene order in pepper and tomato. Proc Natl Acad Sci U S A, 85(17):6419–6423.

36. Bonierbale MW, Plaisted RL, Tanksley SD. 1988. RFLP maps based on a common set of clones reveal modes of chromosomal evolution in potato and tomato. Genetics, 120(4):1095–1103.

37. Wu F, Eannetta NT, Xu Y, Tanksley SD. 2009. A detailed synteny map of the eggplant genome based on conserved ortholog set II (COSII) markers. Theoretical and Applied Genetics, 118(5):927–935.

38. Saei A, Hunter D., Hilario E, David C., Ireland H., Esfandiari A., et al. (2024). De novo Chromosome-level Assembly and Annotation of Petunia hybrida Genome. figshare. Dataset. 10.6084/m9.figshare.26940946

