## Supplementary Figure S1 for "Chromosome-level Genome Assembly and Annotation of *Petunia hybrida*"

**Supplementary Figure S1:** Estimation of genome size and heterozygosity based on 33-mer frequency distribution analysis using the GenomeScope v1. The genome of *P. hybrida* was estimated as 1.15 Gb with heterozygosity of 0.14%.

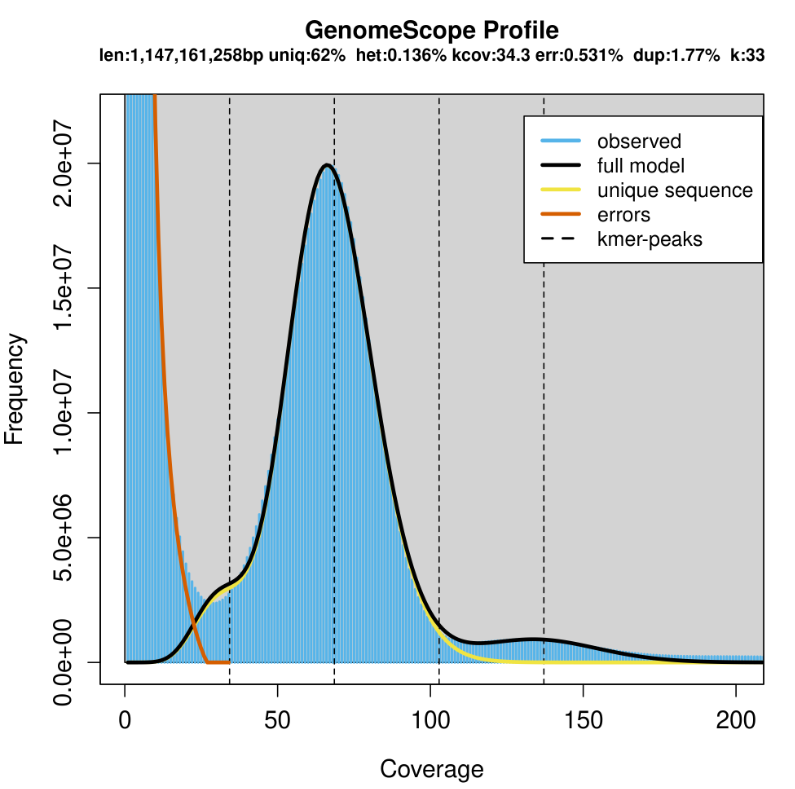
