## Supplementary Figure S3-S4 for "Chromosome-level Genome Assembly and Annotation of *Petunia hybrida*"

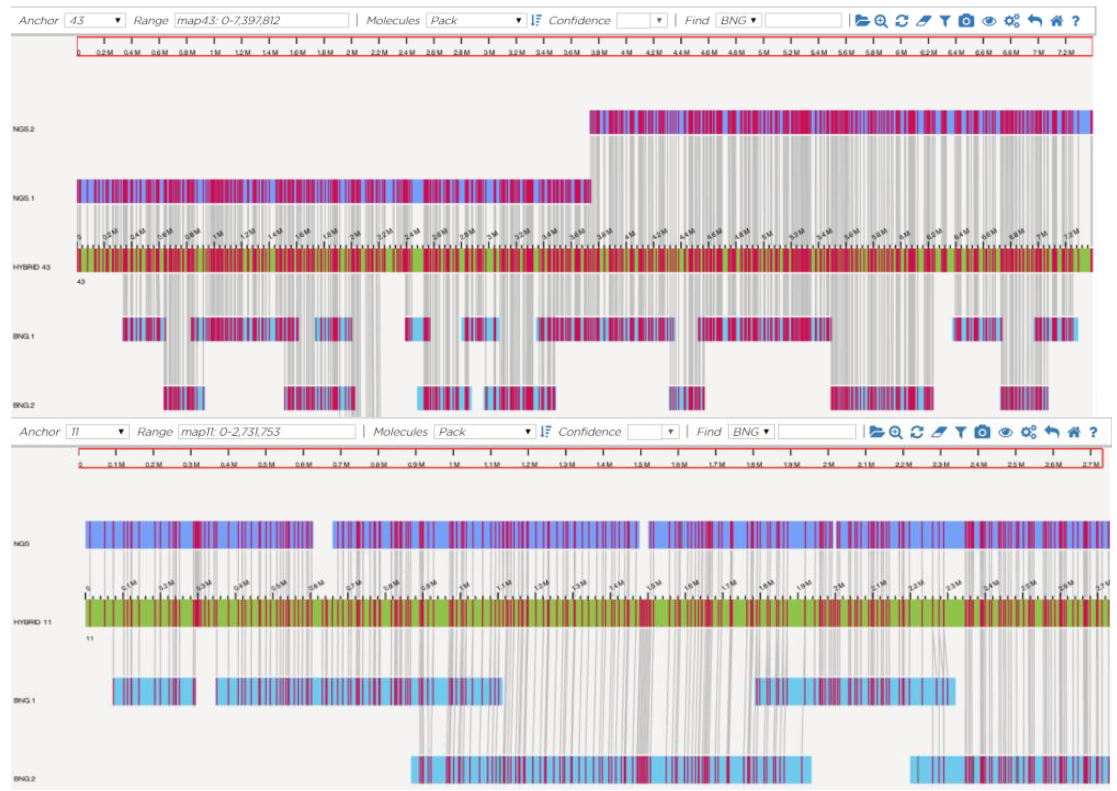

**Supplementary Figure S3:** Visual illustration of how *de novo* assembled optical maps order, orientate, join and confirm sequence integrity of the PacBio contigs. Middle sequence in green is the hybrid scaffold that results from the optical maps (beneath) aligning with each other and with the PacBio contigs (above).

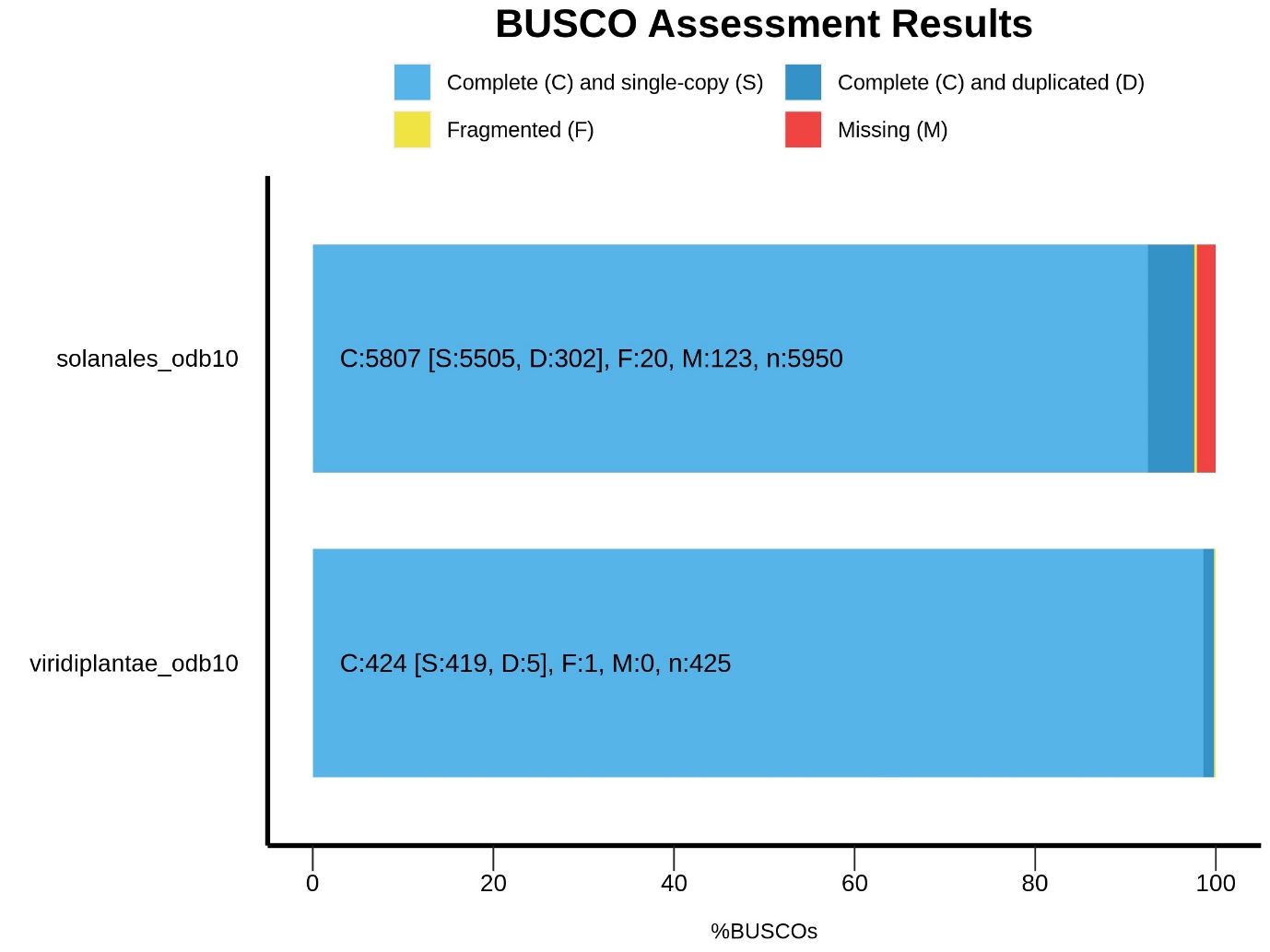

**Supplementary Figure S4.** BUSCO assessment of the *Petunia hybrida* genome completeness using two lineage-specific databases: Viridiplantae_odb10 and Solanales_odb10.
